# The novel, recurrent mutation in the *TOP2A* gene results in the enhanced topoisomerase activity and transcription deregulation in glioblastoma

**DOI:** 10.1101/2020.06.17.158477

**Authors:** Bartlomiej Gielniewski, Katarzyna Poleszak, Adria-Jaume Roura, Paulina Szadkowska, Sylwia K. Krol, Rafal Guzik, Paulina Wiechecka, Marta Maleszewska, Beata Kaza, Andrzej Marchel, Tomasz Czernicki, Andrzej Koziarski, Grzegorz Zielinski, Andrzej Styk, Maciej Kawecki, Cezary Szczylik, Ryszard Czepko, Mariusz Banach, Wojciech Kaspera, Wojciech Szopa, Mateusz Bujko, Bartosz Czapski, Miroslaw Zabek, Ewa Izycka-Swieszewska, Wojciech Kloc, Pawel Nauman, Joanna Cieslewicz, Bartosz Wojtas, Bozena Kaminska

## Abstract

**Background:** High grade gliomas (HGGs) are aggressive, primary brain tumors with poor clinical outcomes. We aim to better understand glioma pathobiology and find potential therapeutic susceptibilities.

**Methods:** We designed a custom panel of 664 cancer- and epigenetics-related genes, and employed targeted next generation sequencing to study the genomic landscape of somatic and germline variants in 182 gliomas of different malignancy grades. mRNA sequencing was performed to detect transcriptomic abnormalities.

**Results:** In addition to known alterations in *TP53*, *IDH1*, *ATRX*, *EGFR* genes found in this cohort, we identified a novel, recurrent mutation in the *TOP2A* gene coding for Topoisomerase 2A occurring only in glioblastomas (GBM, WHO grade IV gliomas). Biochemical assays with recombinant proteins demonstrated stronger DNA binding and DNA supercoil relaxation activities of the variant proteins. GBM patients carrying the mutated *TOP2A* had shorter overall survival than those with the wild type *TOP2A*. Computational analyses of transcriptomic data showed that GBMs with the mutated *TOP2A* have different transcriptomic patterns suggesting higher transcriptomic activity.

**Conclusion:** We identified a novel TOP2A E948Q variant that strongly binds to DNA and is more active than the wild type protein. Our findings suggest that the discovered *TOP2A* variant is gain–of-function mutation.

**Key points:**

- The most frequent genetic alterations in high grade gliomas are reported.
- A new mutation in the *TOP2A* gene was found in 4 patients from Polish population.
- A E948Q substitution changes TOP2A activities towards DNA.
- The recurrent *TOP2A* variant is a gain-of-function mutation.

**Importance of the study:** Glioblastoma is a deadly disease. Despite recent advancements in genomics and innovative targeted therapies, glioblastoma therapy has not shown improvements. Insights into glioblastoma biology may improve diagnosis, prognosis, and treatment prediction, directing to a better outcome. We performed targeted sequencing of 664 cancer genes, and identified a new variant of the *TOP2A* gene encoding topoisomerase 2A in glioblastomas. The TOP2A protein variant shows a higher affinity towards DNA and causes transcriptional alterations, suggesting a higher *de novo* transcription rate.

## Introduction

High-grade gliomas (HGGs) are most frequent, malignant primary brain tumors in adults.^24^ The World Health Organization (WHO) classifies HGGs as WHO grade III and IV gliomas. Among HGGs, the most aggressive is glioblastoma (GBM, a WHO grade IV tumor) which has a high recurrence and mortality rate due to highly infiltrative growth, genetic alterations and therapy resistance. Overall survival of HGG patients ranges from 12-18 months from a time of diagnosis, despite combined treatments encompassing surgical resection, radiotherapy and chemotherapy.^32^ Recent advancements in high-throughput genomic and transcriptomic studies performed by The Cancer Genome Atlas (TCGA) consortium^6^ brought emerging insights into etiology, molecular subtypes and putative therapeutic cues in HGGs. Recurrent somatic alterations in genes such as *TP53*, *PTEN*, *NF1*, *ATRX*, *EGFR*, *PDGFRA* and *IDH1/2* have been reported.^5, 36^ Discovery of the recurrent mutation in the *IDH1* gene^25^ in gliomas and resulting distinct transcriptomic and DNA methylation profiles in *IDH*-mutant and *IDH*-wild type gliomas^7^ improved diagnosis.^20^ Despite advances in understanding glioma genetics, molecularly targeted therapies have, to date, failed in phase III trials in GBM patients. Due to the lack of an effective treatment, even rare but targetable genetic changes are of interest, as they may offer a personalized treatment for selected patients. *BRAF V600E* mutations found in approximately 1% of glioma patients are considered as a therapeutic target in basket trials, including gliomas.^13^ Aurora kinases (AURK) regulate the cell division through regulation of chromosome segregation and their inhibitors have a synergistic or sensitizing effect with chemotherapy drugs, radiotherapy or other targeted drugs in GBMs. Several of AURK inhibitors are currently in clinical trials.^3, 10^

In the current study, we applied targeted next generation sequencing of a custom panel of 664 cancer- and epigenetics-related genes to study the occurrence of somatic and germline variants in 182 high grade gliomas, vast majority of them being glioblastomas. Besides previously described alterations in *TP53*, *IDH1*, *ATRX*, *EGFR* genes, we found several new variants that were predicted to be potential drivers. In four GBM patients from the Polish HGG cohort, we identified a novel, recurrent mutation in the *TOP2A* gene, resulting in a substitution of glutamic acid (E) 948 to glutamine (Q). The *TOP2A* gene encodes Topoisomerase 2A which belongs to a family of Topoisomerases engaged in the maintenance of a higher order DNA structure. These enzymes are implicated in other cellular processes such as chromatin folding and gene expression.^28^ Using computational methods we predicted the consequences of the E948Q substitution on TOP2A and verified them with biochemical assays. Our results show that the E948Q substitution in TOP2A affects its DNA binding and enzymatic activity, which could lead to a “gain-of-function”.

## Materials and Methods

### Tumor samples

Glioma samples and matching blood samples were obtained from the: St. Raphael Hospital, Scanmed, Cracow, Poland; Medical University of Silesia, Sosnowiec, Poland; Medical University of Warsaw, Poland; Military Institute of Medicine, Warsaw, Poland; Copernicus Hospital PL, Gdansk, Poland; Medical University of Gdańsk, Poland; Institute of Psychiatry and Neurology, Warsaw, Poland; Mazovian Brodnowski Hospital, Warsaw, Poland; Maria Sklodowska-Curie National Research Institute of Oncology, Warsaw, Poland; Canadian Human Tissue Bank. Each patient provided a written consent for the use of tumor tissues. All procedures were in accordance with the ethical standards of the institutional and/or national research committees, with the 1964 Helsinki declaration and its later amendments. In the analysis we used 207 glioma samples of WHO II, III and IV grades. For we performed targeted sequencing of DNA from 182 samples; in case of 67 samples, DNA isolated from blood was sequenced as a reference for an analysis of somatic mutations. For the samples provided by Canadian Human Tissue Bank (GL162-GL182) we do not have access to clinical data, excluding diagnosis. Detailed description of the patient cohort is presented in Table 1.

### DNA/RNA Isolation

Total DNA and RNA were extracted from fresh frozen tissue samples using Trizol Reagent (Thermo Fischer Scientific, Waltham, MA, USA), following manufacturer’s protocol. DNA from blood samples was isolated using QIAamp DNA Blood Mini Kit (Qiagen, Hilden, Germany), following manufacturer’s protocol. Quality and quantity of nucleic acids were determined by Nanodrop (Thermo Fisher Scientific, Waltham, MA, USA) and Agilent Bioanalyzer (Agilent Technologies, Santa Clara, CA, USA).

### Design of Targeted Cancer-Related Gene Enrichment Panel

We used SeqCap EZ Custom Enrichment Kit (Roche, Basel, Switzerland, cat. no 06266339001), which is an exome enrichment design that targets the latest genomic annotation GRCh38/hg38. This SeqCap EZ Choice Library covers the exomic regions of 664 genes frequently mutated in cancer. Detailed description of the method and gene panel is presented in Wojtas et al. 2019.^39^

### DNA and RNA Sequencing

DNA isolated from tumor samples was processed for library preparation according to a NimbleGen SeqCap EZ Library SR (v 4.2) user guide. Libraries were prepared from 1 μg of total DNA. Initial step in the procedure was fragmentation of the material to obtain 180-220 bp DNA fragments. Next ends of these fragments were repaired and A base was added to 3’ end. Then indexed adapters were ligated and libraries were amplified. Obtained libraries were mixed eximolar and special oligonucleotide probes were added to enrich the libraries with earlier fragments of interest. After 72 h incubation of the mixture pooled libraries were purified using special beads and amplified. Quality control of obtained libraries was done using Agilent Bioanalyzer with High Sensitivity DNA Kit (Agilent Technologies, Palo Alto, CA, USA). Quantification of the libraries was done using Quantus Fluorometer and QuantiFluor Double Stranded DNA System (Promega, Madison, Wisconsin, USA). Libraries were run in the rapid run flow cell and were paired-end sequenced (2×76bp) on HiSeq 1500 (Illumina, San Diego, CA 92122 USA).

mRNA sequencing libraries were prepared using KAPA Stranded mRNAseq Kit (Roche, Basel, Switzerland, cat# 07962207001) according to manufacturer’s protocol. Detailed description is presented in Rajan et al. 2019.^27^

### Data Analysis

#### Somatic and rare germline variant pipeline

Sequencing reads were filtered by trimmomatic program.^2^ Filtered and trimmed reads were mapped to the human genome version hg38 by BWA aligner^16^ (http://bio-bwa.sourceforge.net/). Next, sample read pileup was prepared by samtools mpileup and then variants were called by bcftools and filtered by varfilter from vcfutils to include just variants with coverage higher than 12 reads per position and with mapping quality higher than Q35. Next, variants were filtered to include only those with a variant allele frequency (VAF) higher than 20%, meaning that more than 20% of reads had to support a variant. A separate analysis on the same sample was performed with a Scalpel software^11^ to enrich for discovery of short deletions and insertions. Variants obtained with a Scalpel were filtered to include only variants with VAF>20%. Variants obtained from bcftools and a Scalpel analysis were merged, and processed to maf format using vcf2maf program. Vcf2maf annotates genetic variants to existing databases by Variant Effect Predictor (VEP) program.^23^ Subsequently, a maf object was filtered by the Minor Allele Frequency (MAF) value, it had to be lower than 0.001 in all populations annotated by VEP to be included in the final analysis, meaning that in any of well-defined populations, including gnomAD, 100 genomes, EXAC it was not found in more than 1 in 1000 individuals. Filtered maf object was subsequently analyzed in maftools R library.^22^ At the end the analyzed glioma cohort included putative somatic and rare germline variants.

#### Somatic variants pipeline

FASTQ files from both tumor and blood DNA samples were mapped to the human genome (hg38) using NextGenMap aligner (http://cibiv.github.io/NextGenMap/). Read duplicates were marked and removed by Picard (https://broadinstitute.github.io/picard/). For somatic calls, a minimum coverage of 10 reads was established for both normal and tumor samples. Additionally, variants with strand supporting reads bias were removed. Only coding variants with damaging predicted SIFT values (>0.05) were selected.

ProcessSomatic method from VarScan2^15^ was applied to extract high confidence somatic calls based on variant allele frequency and Fisher’s exact test p-value. The final subset of variants was annotated with Annovar (http://annovar.openbioinformatics.org/en/latest/) employing the latest versions of databases (refGene, clinvar, cosmic, avsnp150 and dbnsfp30a). Finally, maftools R library^22^ was used to analyze the resulting somatic variants.

To test overlap between a somatic analysis pipeline (including blood DNA as a reference) and somatic and rare germ-line variants pipelines, we compared 50 samples that were common in two analyses. A number of Missense Mutations, Frame Shift Del, Frame Shift Ins and Nonsense mutation obtained was compared between these two pipelines.

#### RNA sequencing analysis

RNA sequencing reads were aligned to the human genome by gap-aware algorithm - STAR and gene counts were obtained by featurecounts.^18^ Aligned reads (bam format) were processed by RseQC^37^ program to evaluate the quality of obtained results, but also to estimate the proportion of reads mapping to protein coding exons, 5’-UTRs, 3’-UTRs and introns. From the gene counts subsequent statistical analysis was performed in R using NOIseq^34^ and clusterProfiler^40^ libraries.

#### Computational prediction of the *TOP2A* mutant effect

To predict the consequences of E948Q substitution on the TOP2A, we used the crystal structure of the human TOP2A in a complex with DNA (PDB code: 4FM9). Protein structures were visualized and an electrostatic potential was calculated using PyMOL.

#### Cloning and mutagenesis

The *TOP2A* gene (NCBI nucleotide accession number (AC) – NM_001067, protein – NP_001058.2) cloned into pcDNA3.1+/C-(K)-DYK vector (Clone ID OHu21029D) was purchased from Genscript. The cDNAs were cloned into a bacterial expression vector. The *TOP2A* fragment corresponding to amino acid residues 890-996 was amplified and cloned into pET28a vector as a BamHI - EcoRI fragment, resulting in a construct expressing His6 tag fused to N-terminus of TOP2A 890-996 aa. The *TOP2A* fragment corresponding to amino acid residues 431-1193 was amplified and cloned into a pET28a vector as a SalI - NotI fragment, resulting in a construct expressing His6 tag fused to N-terminus of TOP2A 431-1193 aa. Site-directed mutagenesis of *TOP2A* gene was performed using a PCR-based technique and a site mutagenesis kit. All resulting constructs were sequenced using Sanger sequencing.

#### Protein expression and purification

Proteins were expressed from bacterial plasmid DNA carrying *TOP2A* in *E. coli* Rosetta (DE3) strain. Cells were grown in LB media with appropriate antibiotics at 37°C. When the bacteria reached OD_600_ 0.8, protein expression was induced by addition of 1 mM IPTG. The induction of TOP2A 890-996 aa was carried out at 37°C for 4 h. One hour before the induction of TOP2A 431-1193 aa the temperature was changed to 37°C and the induction was carried out overnight. Cells were harvested by centrifugation (4000 g for 5 min, 4°C) and pelleted. Proteins were purified using the HisPur Ni-NTA Magnetic beads (Thermo Scientific). The pellet was resuspended in binding buffer (10 mM Na_2_HPO_4_, 1,8 mM KH2PO4, 2,7 mM KCl, 300 mM NaCl, 10 mM imidazole, pH 8.0, 10 % (v/v) glycerol, 10 mM 2-Mercaptoethanol (BME), 1 mM PMSF) and lysed using Bioruptor Plus sonicator (Diagenode). Protein purification was carried out at 4°C. Proteins from the clarified lysates were bound to Ni-NTA resins, the beads were washed with binding buffer, wash buffer 1 (binding buffer with 2 M NaCl) and wash buffer 2 (binding buffer with 20 mM imidazole). The proteins were eluted with elution buffer (binding buffer with 250 mM imidazole, pH 8.0). Next, the buffer was exchanged to (phosphate-buffered saline, 10 % (v/v) glycerol, 10 mM BME) using G25 Sephadex resin (Thermo Scientific) and Spin-X Centrifuge Tube (Costar). Final protein samples of TOP2A 890-996 aa were analyzed using 15% SDS-PAGE and TOP2A 431-1193 aa samples in 6% SDS-PAGE. Homogeneity of the proteins was approximately 90%. Protein concentrations were estimated by measuring the absorbance at 280 nm.

#### Electrophoretic mobility shift assay (EMSA)

DNA binding assay was performed using the LightShift Chemiluminescent EMSA Kit (Thermo Scientific Cat # 20148) according to manufacturer’s instructions. Binding reactions contained 20 pM biotin-labeled oligonucleotide 60 bp end-labeled duplex from LightShift Chemiluminescent EMSA Kit and 4.25 μM TOP2A 431-1193 aa WT or E948Q variant proteins, in a 10 mM Tris pH 7.5 buffer with 50 mM KCl, 1 mM DTT, 5 mM MgCl_2_ in 30 μl. The reaction mixtures were incubated for 30 min at room temperature, and subjected to electrophoresis (100 V, 8°C) on 6% polyacrylamide gels with 10% glycerol and Tris–borate–EDTA buffer. Then, complexes were transferred onto a 0.45 μm Biodyne nylon membrane (Thermo Scientific Cat # 77016) in a Tris– borate–EDTA buffer and detected by chemiluminescence using a Chemidoc camera (Bio-Rad).

#### Supercoil relaxation assay

The supercoil relaxation reaction was conducted at 37°C for 30 min in a buffer containing 20 mM Tris pH 8.0, 100 mM KCl, 0.1 mM EDTA, 10 mM MgCl2, 1 mM adenosinetriphosphate (ATP), 30 μg/ml bovine serum albumin (BSA), 200 ng supercoiled (SC) pET28a plasmid and 0.1 or 0.2 μM WT or E948Q variant TOP2A fragments. Fragments corresponded to 890-996 and 431-1193 aa of the TOP2A. The resultant reaction mixtures were resolved by electrophoresis on 0.7% agarose gel followed by staining with SimplySafe dye (EURX Cat # E4600-01) and visualization using Chemidoc camera (Bio-Rad).

## Results

### Identification of somatic and rare germline variants in gliomas

The study group contained 207 glioma specimens: 21 were from the Canadian Brain Tumor Bank and 186 were from the Polish glioma cohort. Majority of tumors (182) were high grade gliomas (WHO grades III and IV) and 74% were glioblastomas. The clinical characteristics of patients are summarized in the Supplementary Table 1 and Supplementary Fig.1. We searched for somatic and rare germline variants. Genetic variants databases have significantly improved in the last few years that allows to identify variants occurring with a very low frequency in existing databases (based on EXAC, TOPMED, gnomAD and 1000 genomes projects), even in the absence of a reference blood DNA. As the identified germline variants were exceptionally rare in the general population (threshold of MAF<0.001 in all external databases), it is likely that these variants are pathogenic.

Our approach to find potential new oncogenic drivers was based on OncodriveCLUST spatial clustering method.^33^ OncodriveCLUST estimates a background model from coding-silent mutations and tests protein domains containing clusters of missense mutations that are likely to alter a protein structure. The results of the targeted sequencing indicate a high frequency of altered genes (Supplementary Fig. 2A), and are in agreement with previous findings on the landscape of pathogenic genetic changes in gliomas.^30^ Annotations of variants to genes revealed that top 10 of the most frequently altered genes have genetic changes in more than 80% of samples (Supplementary Fig. 2A). The most altered gene was *TP53*, followed by *IDH1*, *PTEN*, *ATRX* and *EGFR* (Supplementary Fig. 2A). Other genes that were found to be frequently altered included *KDM6B*, *ABL1*, *ARID1A*, *AR* and *NCOA6* (Supplementary Fig. 2A). One of the reasons, why these changes have not been noticed previously, is the fact that these changes are most likely rare germline alterations, and represent mostly in-frame insertions (*KDM6B*) or in-frame deletions (*ABL1*, *ARID1A*, *AR, NCOA6)*. These genetic alterations were found at a very low frequency in the general population (MAF<0.001), and would result in insertion or deletion one or more amino acids to the protein, therefore we consider them potentially pathogenic.

The highest number of genetic alterations occur in genes coding for receptor tyrosine kinases (RTK) of RAS pathway (RTK-RAS), less frequently in genes coding for proteins from NOTCH, WNT and PI3K pathways (Supplementary Fig. 2B). More than 30 genes from the RTK-RAS pathway were found to be altered (Supplementary Fig. 2C). In this cohort, the *ABL1* gene was found to be altered more frequently than the *EGFR* gene (Supplementary Fig. 2C). Most frequently mutated genes encoding specific protein domains from the PFAM database are: the TP53 domain, the PTZ00435 domain related to *IDH1/2* mutations, conserved domains of protein kinases (PKc_like, S_TKc) and kinase phosphatases (PTPc) (Supplementary Fig. 2D).

The results show the *IDH1* gene as a top scoring hotspot, but among identified genes we found also genes such as *FOXO3*, *PDE4DIP*, *HOXA11, ABL1* and *RECQL4* that were not previously reported as mutated in HGGs (Fig.1A, B). When the identified genes were compared to the results obtained in the TCGA project, most of the genes (shown in bold in the Fig.1B) were found to be potentially new oncogenic genes.

**Fig. 1.**
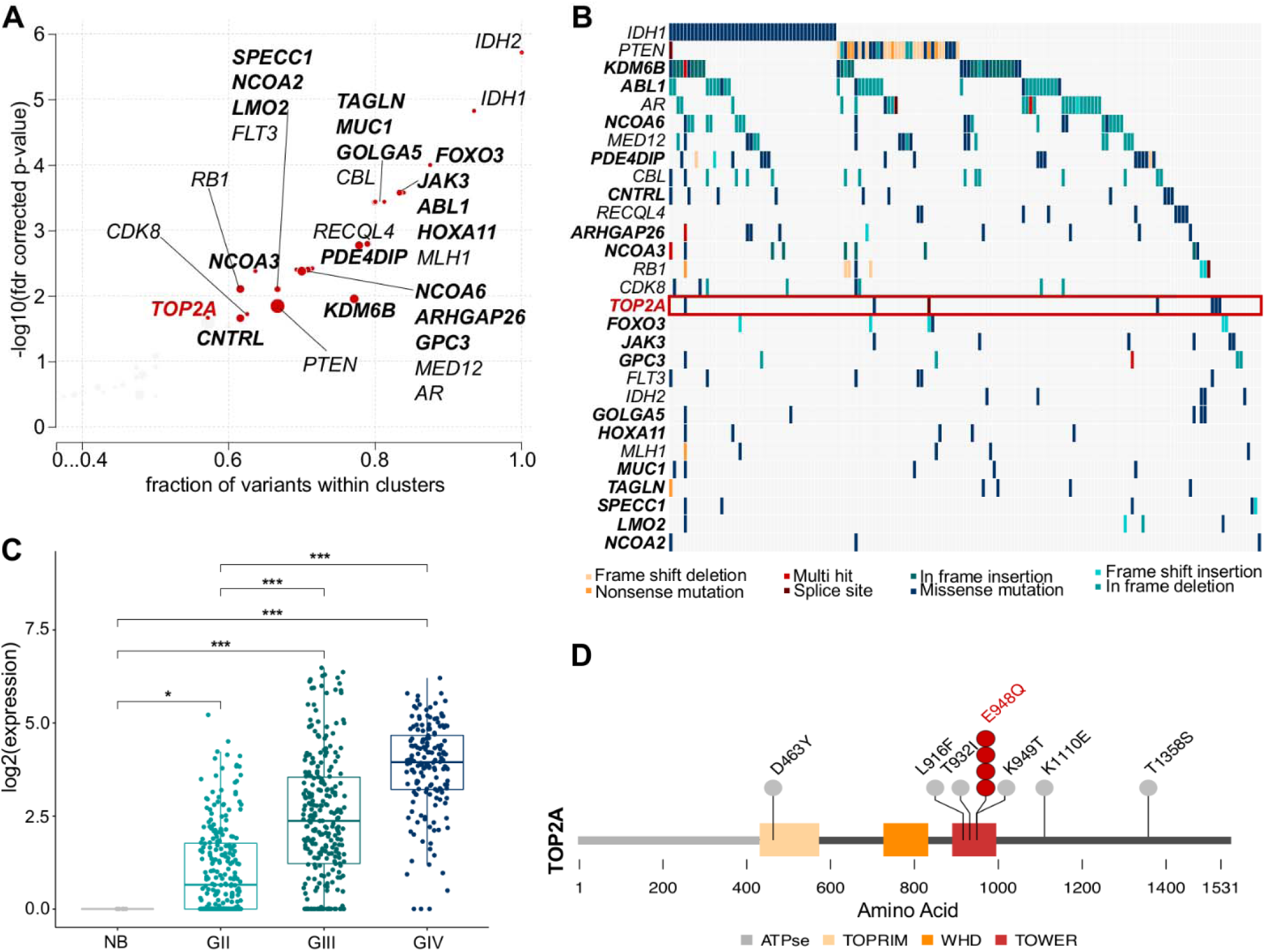
Targeted sequencing identified frequent mutations in high grade gliomas and revealed novel mutations in the *TOP2A* gene. A – Plot showing potential oncogenic drivers (OncodriveCLUST function) in the cohort of high grade gliomas (HGGs), red dots represent potential drivers, with FDR-corrected OncodriveCLUST P value<0.025; size of the dot corresponds to a number of the alterations that were present in the cluster. **A, B**-Genes marked in bold are new potential oncodrivers in HGGs, as these genes were not found to be mutated in TCGA LGG/GBM datasets. **B** – Oncostrip plot showing genes discovered in the OncodriveCLUST analysis, each column corresponds to a glioma patient, while each row corresponds to one altered gene. **C** – *TOP2A* expression in normal brain (NB) and WHO glioma grades (GII, GIII and GIV) in the TCGA dataset. P-values based on Wilcoxon signed-rank test were obtained (* P < 0.05, ** P < 0.01 and *** P < 0.001). **D** – TOP2A lolliplot showing the identified substitutions mapped on the protein structure of TOP2A. A number of dots represents the number of patients in which the particular alteration was found.

One of the most interesting genes was *TOP2A,* which was mutated in several GBMs. The TCGA dataset analysis showed the higher expression of *TOP2A* in high grade gliomas, particularly in glioblastomas (Fig. 1C). In total, the *TOP2A* gene harbored 7 different mutations but interestingly, one mutation was found in 4 GBM patients exactly in the same chromosomal position. This recurrent mutation in the *TOP2A* gene was located inside a gene fragment encoding the TOWER domain of TOP2A (Fig. 1D). The presented lolliplot (Fig. 1D) shows all variants found in the *TOP2A* gene, including the ones that did not pass QC analysis (Supplementary Table 2).

Considering a set of the somatic mutations with deleterious consequences, the most frequently altered gene in the cohort was *TP53*, which is mutated in 48% of the patients, followed by *IDH1*, *ATRX*, *EGFR* and *PTEN*, among others (Supplementary Fig. 3). To verify concordance of the somatic and somatic/rare germline pipeline of the variant analysis, we have computed overlap of these results, and found that most of somatic variants are also detected by a somatic/rare germline pipeline (Supplementary Table 3).

### Novel mutations identified in the *TOP2A* gene and their pathological consequences

We found seven SNPs in the *TOP2A* gene in ten glioblastoma samples (Supplementary Table 2). All identified SNPs give rise to missense variants. Transversion of guanine to cytosine at the chromosomal position 17:40400367, which results in the substitution of E948 to Q, was found in four GBM patients. Both the recurrent one and other detected SNPs in the *TOP2A* gene were found in GBM patients from the Polish HGG cohort. The occurrence of the recurrent SNP in the position 17:40400367 has been verified by Sanger sequencing, apart from one sample, which had a low variant allele frequency. One of identified variants was identified as a somatic variant by comparison with a reference blood DNA; the presence of another SNP was not detected in a reference DNA from patient’s daughter. These partial results combined with a low frequency of this variant in the Polish population suggest a somatic mutation.

The *TOP2A* gene encodes Topoisomerase 2A. Topoisomerases are enzymes that relax, loosen and disentangle DNA by transiently cleaving and re-ligating both strands of DNA, which modulates DNA topology.^9, 28^ The novel, recurrent mutation in the *TOP2A* gene results in a substitution of E948 to Q in the Tower domain of the TOP2A.

We studied exclusivity and co-occurrence of *TOP2A* mutations among detected top mutations. *TOP2A* mutations co-occurred with *ARID1A* mutations, encoding a SWI/SNF family helicase, and *KMT2C* (formerly *MLL3*), coding for histone methyltransferase, as well as with *EGFR* and *NOTCH3* mutations (Fig. 2A). One of the *TOP2A* mutated samples (first from the left in Fig. 2A) is a hypermutated sample, some of the other samples have also a high number of genetic alterations when compared to other well-defined clusters, such as *IDH1_TP53-, IDH1-, TP53- and PTEN-*mutated clusters (Fig. 2B). All *TOP2A* mutated samples show a higher number of genetic alterations when compared to other defined clusters of glioma samples (Fig. 2B). Moreover, most of the changes in *TOP2A* mutated samples are C>T transitions, most likely related to deamination of 5-methylcysteine (Fig. 2C). There is a high variability within GBM samples with the *TOP2A* mutated, with approximately a half of *TOP2A* samples having a very high fraction (>80%) of C>T changes, while the other half have an expected fraction (<40%) of C>T changes (Fig. 2C).

**Fig. 2.**
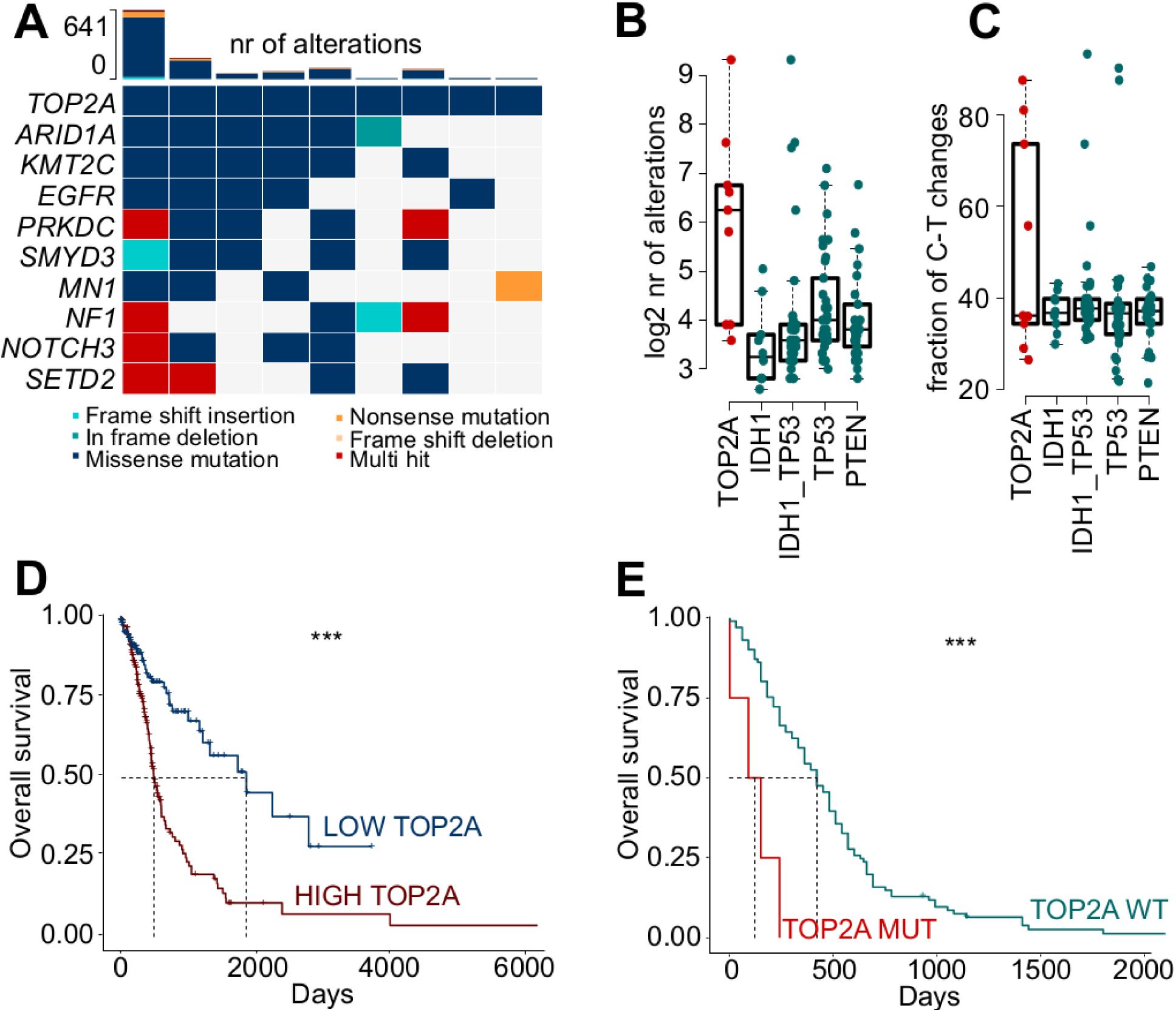
Co-occurrence of genomic alterations of *TOP2A* mutated samples and effects on survival. **A** – Oncoplot showing most frequently altered genes in *TOP2A* mutated HGGs, each column corresponds to a glioma sample, while each row corresponds to one altered gene. Log2 number of genetic alterations in *TOP2A* (**B**) and a fraction of C to T nucleotide changes (**C**) in the *TOP2A* mutated samples versus the samples with other mutations: *IDH1*, *IDH1* and *TP53*, *TP53* and *PTEN*. Sample assignment to each of described groups is explained in the Supp. Fig. 4. **D**-Kaplan-Meier overall survival curve for patients with high or low *TOP2A* expression in the TCGA dataset. Patients alive at the time of the analysis were censored at the time they were followed up. The median overall survival was ~500 days in the HIGH *TOP2A* mRNA expression group and ~2000 days in the LOW *TOP2A* mRNA expression group. P-values based on Log Rank Test were obtained. **E** – Kaplan-Meier overall survival curve in this cohort. The median overall survival was ~150 days in the HGG patients with the *TOP2A* mutated (TOP2A MUT (E948Q) and ~400 days in the patients with wild-type *TOP2A* (TOP2A WT). P-values based on Log Rank Test were calculated.

We sought to determine if expression levels of *TOP2A* or an occurrence of the mutated *TOP2A* version may affect patient’s survival. Using RNA sequencing (RNA-seq) data from HGGs in the TCGA dataset, we demonstrate that patients with the high *TOP2A* levels had shorter survival than other HGG patients (p=0.00027) (Fig. 2D). Survival analysis of patients harboring the recurrent *TOP2A* mutation showed that three GBM patients with the mutated *TOP2A* (one patient died after surgery) survived 5, 3 and 8 months from the time of diagnosis, respectively. Although, a number of patients was low, the data indicate that the GBM patients harboring this *TOP2A* variant had a shorter overall survival compared to other GBM patients (Fig. 2E). This observation suggests a negative effect of mutations in the *TOP2A* gene on the overall survival of GBM patients.

### The TOP2A E948Q substitution may affect protein-DNA interactions

To predict a consequence of E948Q substitution on the TOP2A, we used the crystal structure of the human TOP2A in a complex with DNA (PDB code: 4FM9^38^ (Fig.3A, B). Amino acid residue E948 is located in the Tower domain of the TOP2A protein (Fig. 3C), which with other domains (TOPRIM and WHD) form the DNA-Gate involved in DNA binding.^1^ In the crystal structure of the human TOP2A in a complex with DNA E948 is in a close proximity to DNA (Fig. 3B). The replacement of a negatively charged E residue to a polar Q residue changes an electrostatic potential of the protein (Fig. 3C). Therefore, we predict that this change would affect protein-DNA interactions, and the TOP2A E948Q variant may have an increased affinity towards DNA. Some of the other discovered *TOP2A* mutations may also have the effect on TOP2A binding to DNA, as described in Supplementary Table 4.

**Fig. 3.**
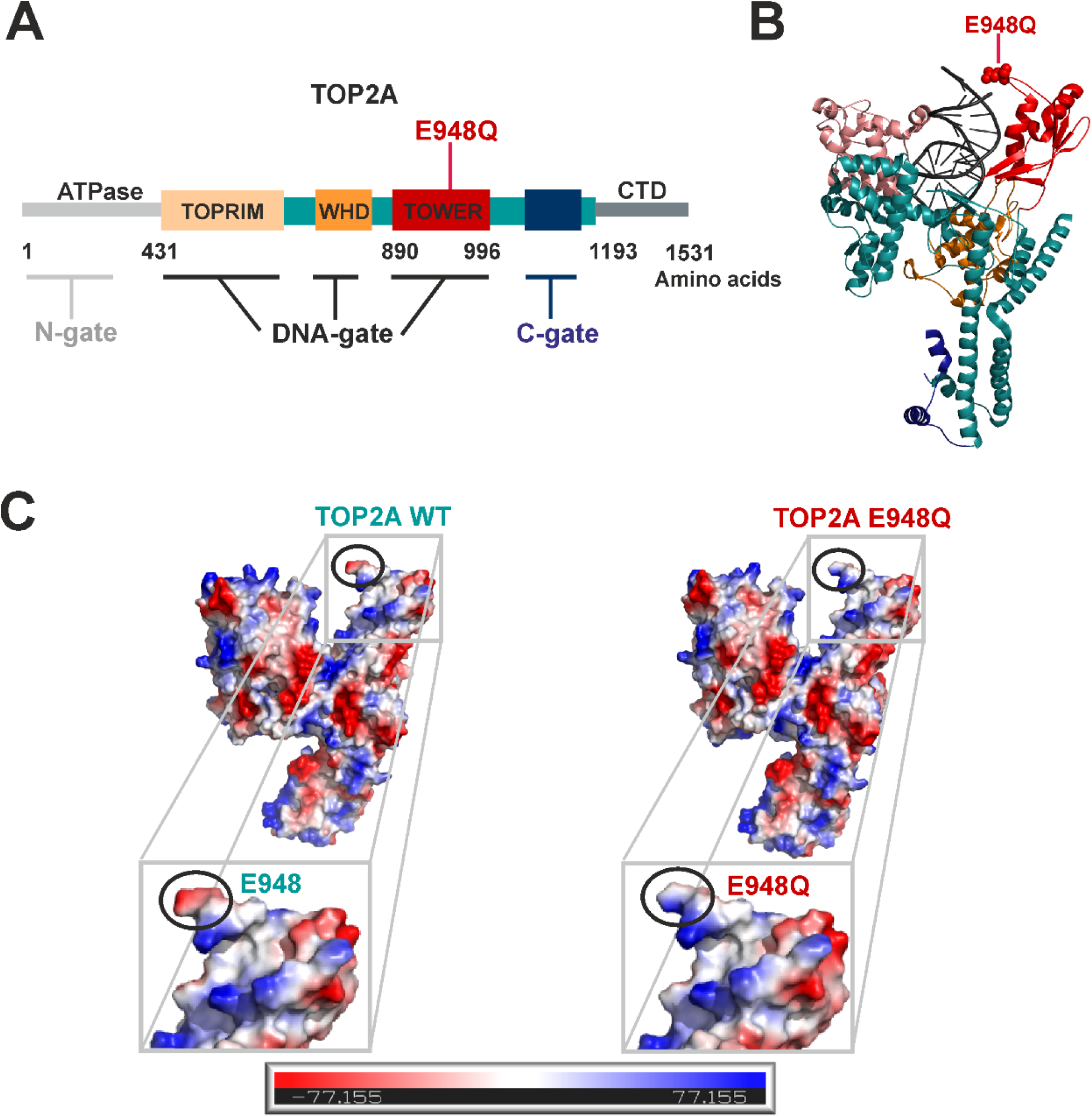
Substitution of E948Q to Q changes the electostatic potential of TOP2A. **A** – Schematic representation of a TOP2A domain structure. Functional regions are colored and labeled: an ATPase domain, a TOPRIM - Mg2+ ion-binding topoisomerase/primase fold, a WHD – winged helix domain, a TOWER, CTD – C-terminal domain. The three dimerization regions are marked: N-gate, DNA-gate and C-gate. **B** – Secondary structure representation of TOP2A 431-1193 bound to DNA (crystal structure PDB code: 4FM9). For simplification only a TOP2A protomer is shown. An amino acid residue E948, which is substituted to Q in a TOP2A variant is represented as spheres on side chains. **C** – Electrostatic surface potential of TOP2A 431-1193 WT and E948Q variant. Electrostatic potential was calculated using a PyMOL program.

### The TOP2A E948Q substitution increases the enzyme binding to DNA and its enzymatic activity

To analyze the effect of the TOP2A E948Q substitution on an enzymatic activity, we produced and purified recombinant wild-type (WT) and variant TOP2A proteins as His6 tag-fusion proteins (Fig. 4A), and performed biochemical tests. We compared a DNA binding activity of WT and E948Q TOP2A fragments using an electrophoretic mobility shift assay (EMSA) and purified proteins. We tested binding of a 890-996 aa fragment of TOP2A comprising only the Tower domain and a 431-1193 aa fragment of TOP2A, which includes the entire DNA-Gate domain (see Materials and Methods). The representative EMSA gel shows that the TOP2A variant E948Q binds to DNA more strongly (Fig. 4B). The lower panel shows the results of densitometric assessment of shifted bands from two experiments.

**Fig. 4.**
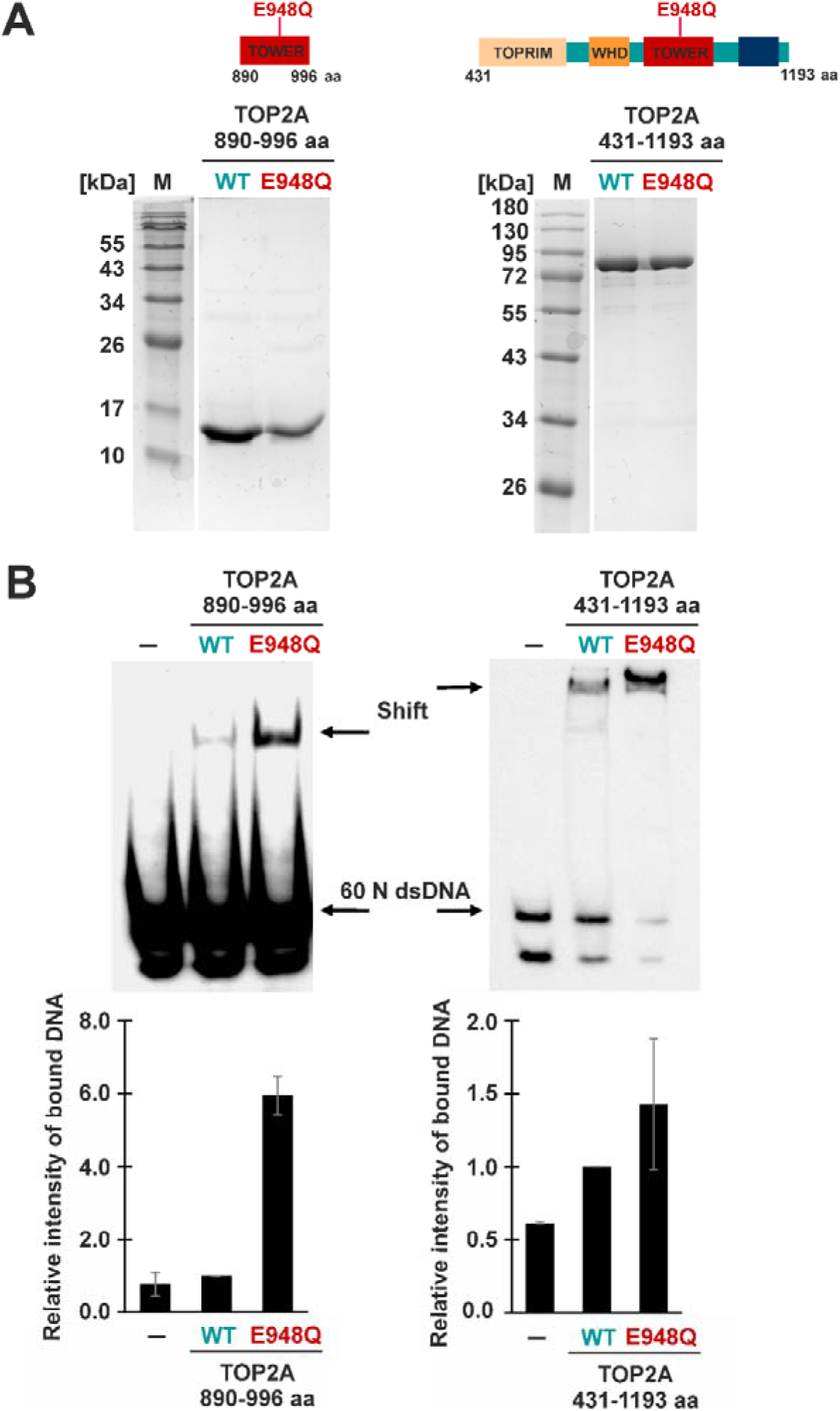
TOP2A E948Q protein exhibits a higher DNA binding capacity than WT. **A** – Purified TOP2A proteins. Schematic representation of purified proteins is placed at the top of the figure: TOP2A 890-996 aa (only the TOWER domain) and TOP2A 431-1193 aa (the DNA-gate). Amino acid residue E948, which is substituted to Q in the TOP2A, is labeled in the scheme. SDS-PAGE of TOP2A purified proteins. Purified proteins were resolved in a 6% and 15% SDS–PAGE and stained with Coomassie brilliant blue. **B** – DNA-binding activity of TOP2A WT and E948Q. EMSA was performed using the LightShift Chemiluminescent EMSA Kit. The binding reactions were resolved in 6% polyacrylamide gels, transferred onto a membrane and detected by chemiluminescence. Binding activities are expressed in relation to that of TOP2A WT (which was set to 1), and are presented as mean ± standard deviations (error bars) that were calculated from two independent measurements. Non-specific binding was determined in a reaction without a protein and was subtracted from the results.

Subsequently, we analyzed the effects of the TOP2A E948Q substitution on a catalytic activity of the protein. We determined a supercoil DNA relaxation activity of WT and TOP2A E948Q proteins. Disappearance of the lower band representing a supercoiled plasmid DNA (SC bands), and appearance of a higher band representing an untwisted plasmid DNA (OC) shows differences in the enzymatic activity of TOP2A proteins. The lower panel shows the results of densitometric assessment of shifted bands from these experiments. We conclude that the E948Q TOP2A protein is functional, and might have a higher activity than the WT protein (Fig. 5).

**Fig. 5.**
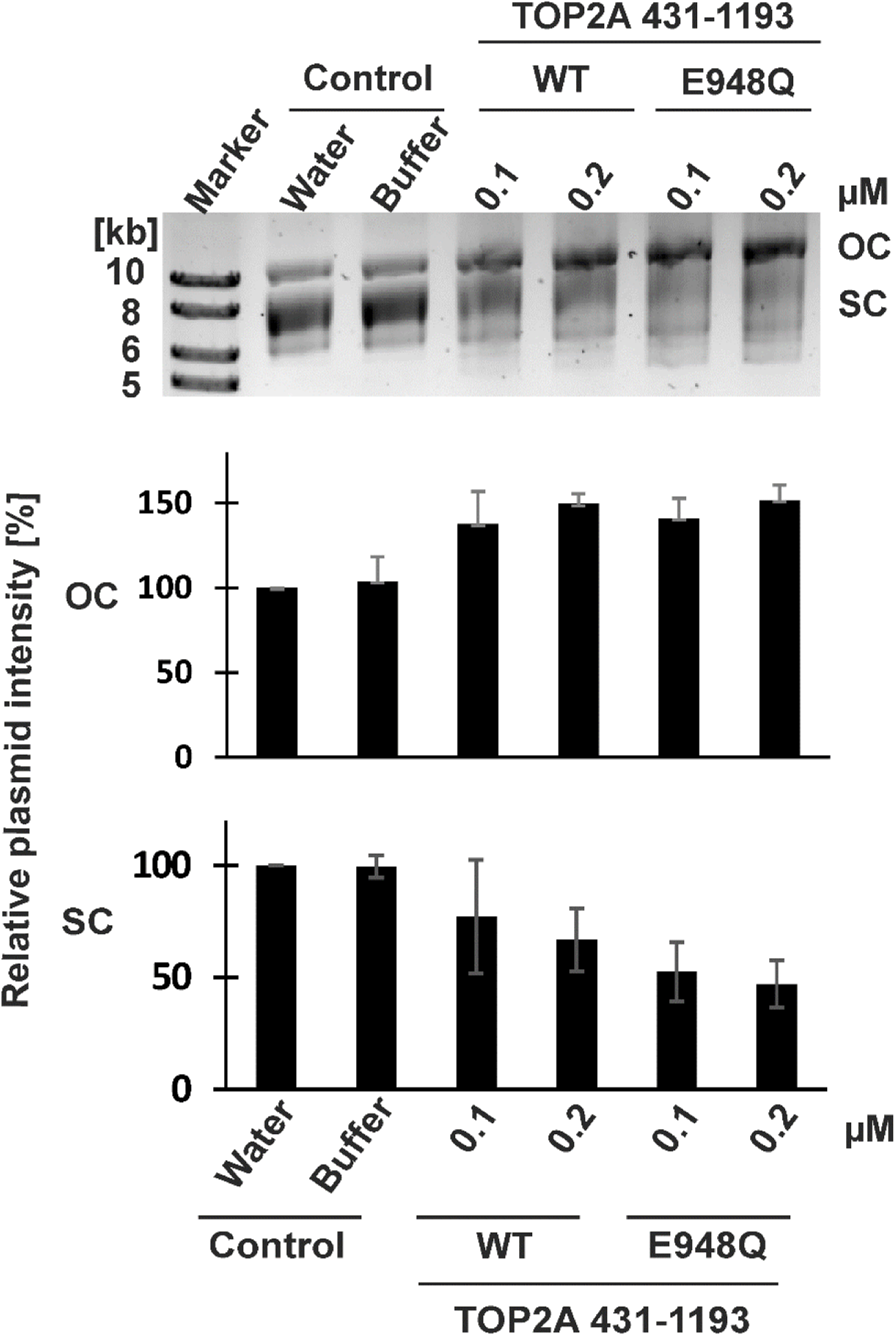
Activity of TOP2A WT and E948Q variant. Supercoil relaxation assay of a plasmid DNA induced by TOP2A 431-1193 WT or E948Q proteins. The reaction mixtures were resolved by electrophoresis on 0.7% agarose gel followed by staining with SimplySafe dye and visualization using a Chemidoc camera. The protein activities were measured by its ability to unwind a supercoiled plasmid DNA (SC) to an open circular DNA (OC). The intensities of SC and OC plasmid DNA were measured in relation to a control DNA without an added protein (which was set to 100%) and are presented as means ± standard deviations (error bars) that were calculated from three independent measurements.

### Transcriptomic alterations in glioblastomas with the mutated *TOP2A* point to dysfunction of splicing

RNA sequencing was performed on 105 samples of low (WHO grade II) and high-grade gliomas (WHO grades III and IV). We searched for any peculiarities of transcriptional patterns in the wild type and mutated *TOP2A* gliomas. Among the analyzed samples were 7 GBM samples with mutations in the *TOP2A* gene, including 4 samples with a recurrent mutation at position 17: 40400367. Transcriptional patterns of GBMs with mutations in the *TOP2A* gene were compared to samples with mutations in the *IDH1, ATRX, TP53 and PTEN* genes.

We found that higher percentages of the detected sequencing reads are mapped in non-coding (intron) regions in GBMs with the mutated the *TOP2A* gene when compared to gliomas with the *IDH1, ATRX, TP53,* and *PTEN* mutations (Fig. 6A). This observation suggests a higher level of transcription in samples with the mutated *TOP2A*. The higher content of intron regions in the RNA sequencing data may indicate a higher level of *de novo* transcription or deficiency in mRNA splicing (Fig. 6A). An analysis of the content of G-C pairs in the coding sequences of the expressed genes in gliomas with the mutated *TOP2A* versus other samples was performed (Fig. 6B). In GBMs with the *TOP2A* mutation, the genes with the coding sequences rich in A-T pairs were more often expressed, while the genes with coding sequences rich in G-C pairs were less frequently present in these samples (Fig. 6B).

**Fig. 6.**
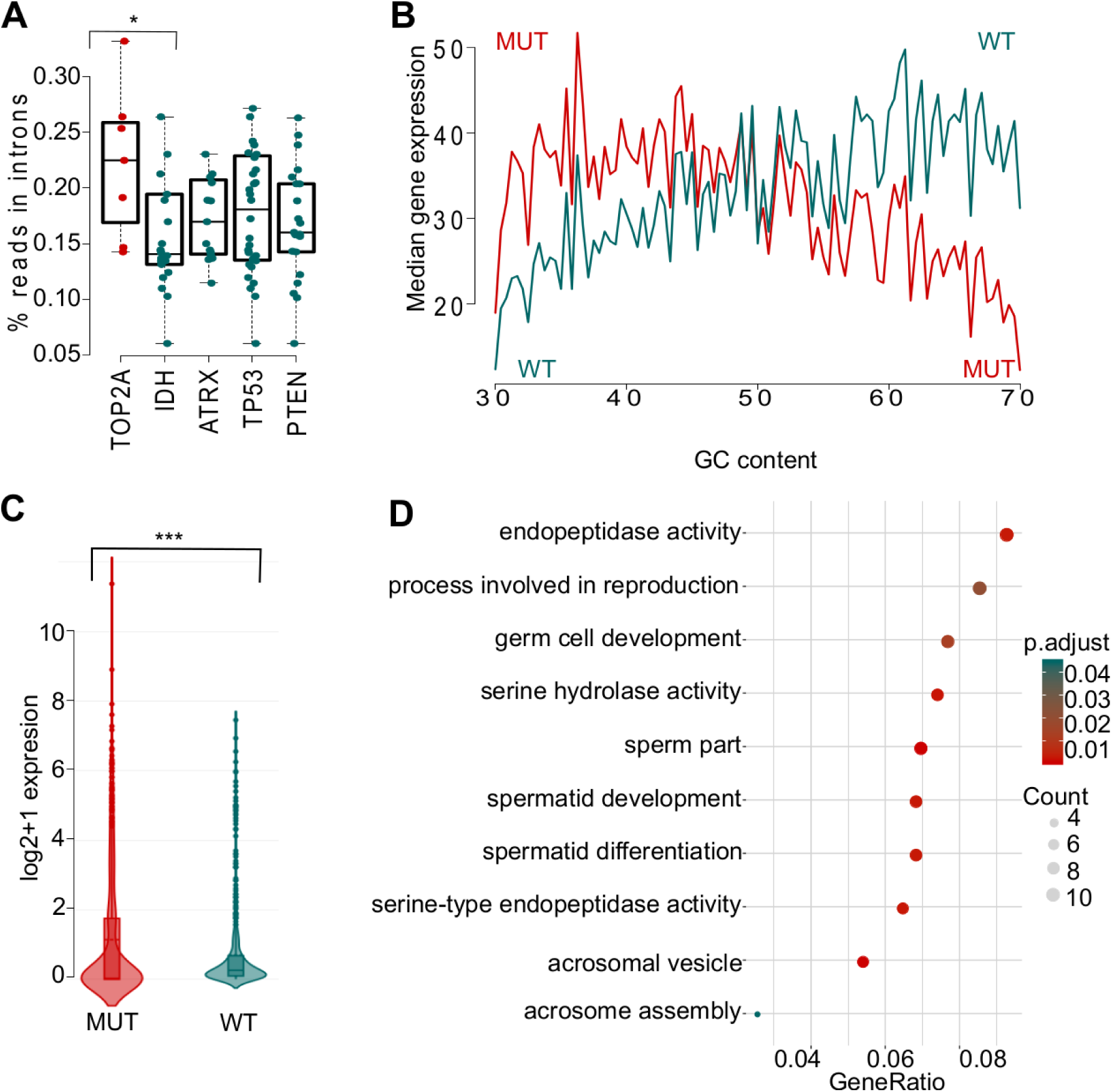
Transcriptomic alterations in *TOP2A* mutated HGGs. **A** – Fractions of RNA-seq reads mapped to intron regions. **B**-GC content profile of gene expression in *TOP2* mutant (MUT) and wild type (WT) HGGs. **C** – Differences in expression of genes that are not expressed in the normal brain in *TOP2A* MUT and WT gliomas. Genes were preselected by filtering out all the genes that were expressed in any of the cells from Stanford Human Brain RNAseq containing data from sorted healthy brain cells. Additionally, all genes expressed in normal brain samples were filtered out. **D** – Gene Ontology terms show functional annotations of genes not expressed in the normal brain that became activated in *TOP2A* mutant HGGs. P-values for plots A based on Wilcoxon signed-rank test, for the panel C on Kolmogonorov-Smirnov test (* P < 0.05, ** P < 0.01 and *** P < 0.001).

Additionally, we compared the expression of a set of genes (that are not expressed in the normal brain) in *TOP2A*-wild type (WT) and *TOP2A* mutated GBMs. RNA-seq data were filtered to remove all brain-specific genes, by filtering out genes expressed in the brain uploaded from the Stanford Brain RNA-Seq database and our data on human normal brain samples. Expression of non-brain specific genes was higher in the *TOP2A* mutated samples than in *TOP2A* WT glioma samples (Fig. 6C). We performed a Gene Ontology (GO) analysis of differentially expressed, non-brain specific genes, and the results showed over-representation of pathways involved in an endopeptidase activity and spermatid development, suggesting acquisition of some new functionalities related to aberrant differentiation and migratory processes (Fig. 6D).

GO terms analysis of non-brain specific genes that are upregulated in *TOP2A* mutants showed increased expression of genes associated with an endopeptidase activity and germ cells development in samples with the *TOP2A* mutations (Fig.6D). These newly acquired functions of *TOP2A* mutants may contribute to a shorter survival time of *TOP2A* mutated GBM patients.

## Discussion

The targeted NGS-based genomic profiling of 664 cancer-related genes used in the present study was capable of detecting genomic alterations in glioma specimens at high sensitivity and specificity. We identified a number of well-known and frequent alterations in *TP53*, *IDH1*, *ATRX*, *EGFR*, *CDK8, RB1, PTEN* genes previously described in glioblastomas, which validates our approach. However, using OncodriveCLUST^33^ we identified several novel variants that represent potentially deleterious germline, somatic mosaicism or loss-of-heterozygosity variants. Mutations in genes such *FOXO3*, *PDE4DIP*, *HOXA11, ABL1, AKAP9, TOP2A* and *RECQL4* were not previously reported in HGGs (Fig.1A, B). In three HGGs we found a recurrent mutation in the *AKAP9* gene coding for A-kinase anchor protein 9/AKAP9), a scaffold protein which regulates microtubule dynamics and nucleation at the Golgi apparatus.^35^ AKAP9 promotes development and growth of colorectal cancer^12^, and is associated with enhanced risk and metastatic relapses in breast cancer.^14^ The somatic variation in the *AKAP9* gene could have a damaging effect on four different transcripts, and may result in a non-functional or truncated protein.

Among top 50 genes most frequently mutated in this cohort, of particular interest are genetic alterations in the genes coding for epigenetic enzymes and modifiers: *KDM6B, NCOA2*, *NCOA3* and *NCOA6.* KDM6B is critical for maintenance of glioma stem cells that upregulate primitive developmental programs.^19^ KDM6B has been targeted with a specific inhibitor GSK-J4 in acute myeloid leukemia^17^, and GSK J4 is active against native and TMZ-resistant glioblastoma cells.^29^ The presence of mutations in genes encoding three different members of the NCOA (nuclear receptor co-activator) family of co-activators suggests the importance of these co-activators in glioma pathogenesis. NCOA2 is a co-activator of diverse nuclear receptors such as estrogen receptor, retinoic acid receptor/retinoid X receptor, thyroid receptor and vitamin D receptor, and regulates a wide variety of physiological responses.^21^ NCOA6 can associate with three co-activator complexes containing CBP/p300 and Set methyltransferases, and may play a role in transcriptional activation by modulating chromatin structure through histone methylation.^21^ The biological relevance of these genomic alterations in HGGs awaits further investigation.

In current study, we focused on a newly identified, recurrent mutation in the *TOP2A* gene present in four GBM patients from the Polish HGG cohort. *TOP2A* mutations were described in many cancers, including breast and lung cancer^8^, but not in gliomas. As this recurrent mutation in the *TOP2A* was discovered only in GBMs from the Polish HGG cohort (but not in 21 GBMs from the Canadian Brain Tumor Bank), we consider this mutation as a putative founder mutation. While a reference blood DNA was accessible only in case of two patients, the data indicate that this mutation is somatic. Topoisomerases regulate DNA topology by introducing transient single- or double-strand DNA breaks, and may relax and unwind DNA, which is a critical step for proper DNA replication or transcription.^26^ Our computational predictions localized a substitution of E948 in the Tower domain of the TOP2A protein in a close proximity to DNA. The replacement of a negatively charged residue E to a polar residue Q would change an electrostatic potential of the protein. This prediction has been confirmed by comparing DNA binding and enzymatic activities of recombinant WT and E948 TOP2A proteins. As TOP2A is a large protein, we prepared constructs expressing the 890-996 aa N-terminus of TOP2A (comprising only the Tower domain) and the 431-1193 aa N-terminus of TOP2A (encompassing the entire DNA-Gate domain). Biochemical assays showed that the TOP2A E948Q binds to DNA more strongly and has a higher supercoil relaxation activity than WT TOP2A.

The presented evidence suggests that TOP2A mutant acquires new functions and exhibits a gain-of-function phenotype. Global gene expression profiling showed different expression of genes with a high AT content and abundance of mRNAs with more intronic regions in GBMs harboring the mutated *TOP2A* gene in comparison with WT *TOP2A* GBMs. These transcriptomic changes suggest either an increased transcription rate or splicing aberrations. Moreover, GBMs harboring the mutated *TOP2A* showed activation of genes encode proteins with an endopeptidase activity that are implicated in cancer invasion. Acquisition of these properties may lead to an increased migratory and invasive phenotype that leads to poor outcome for the patients. GBM patients with the mutated *TOP2A* showed shorter survival than those with WT *TOP2A*. Interestingly, the analysis of the TCGA dataset shows increased expression of *TOP2A* in high grade gliomas, and a negative correlation between *TOP2A* expression and patient’s survival. Altogether, the results support a notion that a newly discovered mutation in the *TOP2A* gene is a gain-of-function mutation, which results in enhancement of the enzymatic activities and acquisition of new functions leading to disturbed transcription rate and altered transcriptomic profiles.

## Acknowledgements

Studies were supported by the Foundation for Polish Science TEAM-TECH Core Facility project “NGS platform for comprehensive diagnostics and personalized therapy in neuro-oncology”. The use of CePT infrastructure financed by the European Union — The European Regional Development Fund within the Operational Programme “Innovative economy” for 2007-2013 is highly appreciated. We thank all the patients for the consent for the use of their biological material for this research.

