## Supplementary materials for "The novel, recurrent mutation in the *TOP2A* gene results in the enhanced topoisomerase activity and transcription deregulation in glioblastoma"

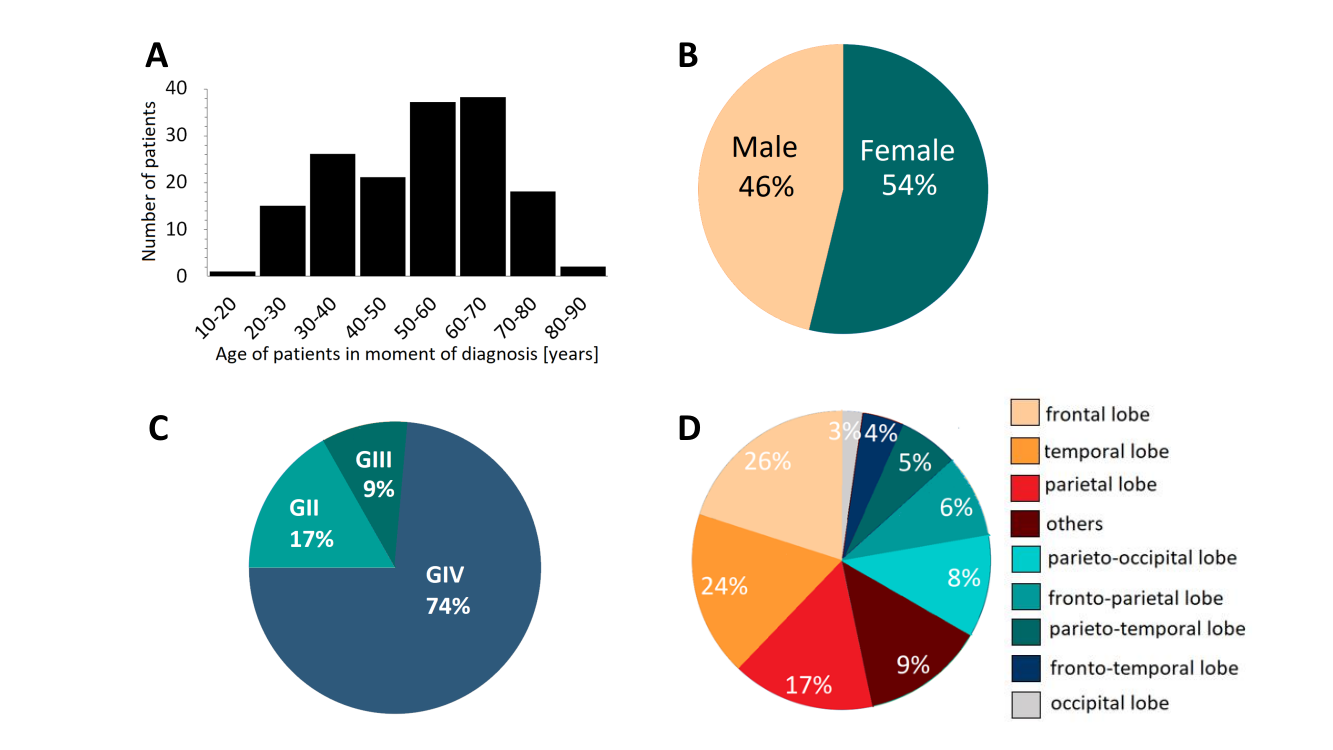

**Supp. Fig. 1**. **Clinical characteristics of patients**. A – patient age distribution. B – patient sex distribution. C – percentages of HGG patients of a specific WHO grade). D – percentages of patients with a specific localization of the tumor in the brain.

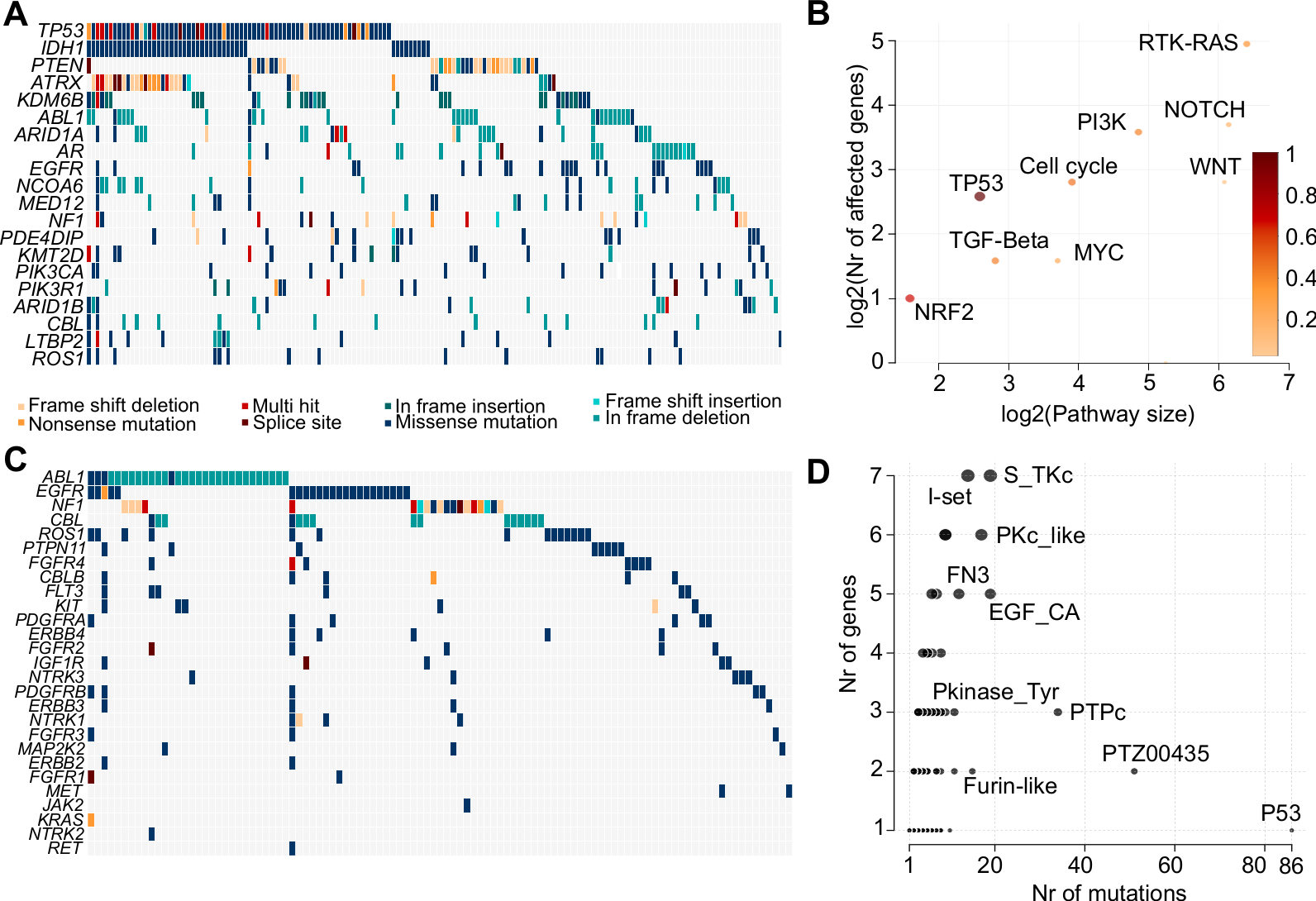

**Supp. Fig. 2.** **The detailed landscape of somatic and rare germ-line variants in high-grade gliomas.** A – Oncostrip showing most frequently altered genes in high-grade gliomas, each column corresponds to a glioma sample, while each row corresponds to one altered gene. B – Plot of oncogenic pathways more frequently altered in the studied cohort of high-grade gliomas. Size and color intensity of dots mark how many genes from a given pathway were mutated in the studied cohort. C – Oncostrip plot of RTK-RAS pathway which emerges as the most frequently altered in this cohort. D – Most commonly mutated pfam domains, size of a dot corresponds to a number of genes with that particular domain being altered in the studied cohort.

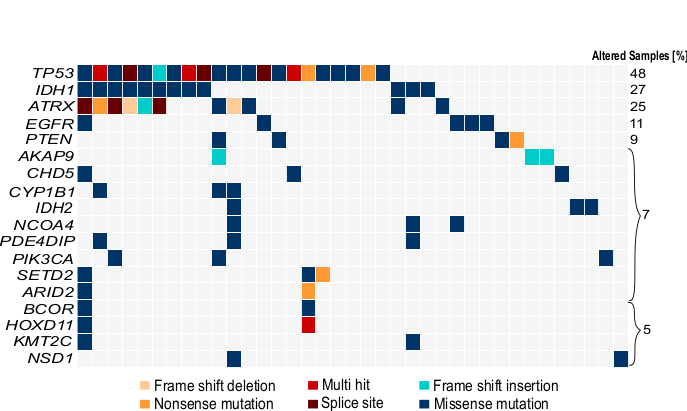

**Supp. Fig. 3.** Oncostrip showing most frequently altered genes in high-grade gliomas, color coded by the specific mutation type.

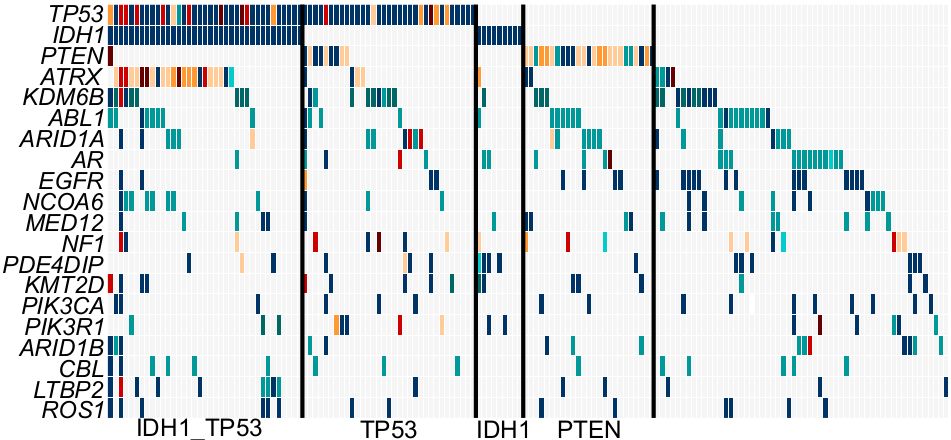

**Supp. Fig. 4.** Oncostrip showing most frequently altered genes in high-grade gliomas, each column corresponds to a glioma sample, while each row corresponds to one altered gene. Vertical lines demark the groups with mutated *IDH1* and*TP53*, *TP53*, *IDH1* and *PTEN* that were used to achieve data presented in the Figs. 3B and C.

**Supplementary Table 1. Clinical characteristics of the tumor samples used in the study**. The following coding was used: "M" - male, "F" - female, "ND" - no data

**Supplementary Table 2. List of genetic changes identified in the TOP2A gene using targeted next generation sequencing of 182 glioma samples.** The table contains the most important information regarding the identified variants

**Supplementary Table 3. Comparison of the number of identified somatic variants using two different data analysis pipelines.** Data analysis pipeline for identifying only somatic variants vs data analysis pipeline for identifying somatic and rare germline variants were compared.

**Supplementary Table 4. The most important features of the amino acid substitutions caused by mutations identified in *TOP2A* gene.** Some of the identified changes may increase or decrease DNA binding.
