## Supplementary Table 3 for "The novel, recurrent mutation in the *TOP2A* gene results in the enhanced topoisomerase activity and transcription deregulation in glioblastoma"

| Type of genetic alteration | Number of mutations detected using analysis for somatic variants | Number of mutations detected using analysis for somatic and rare germline variants | Number of mutations detected in both type of analysis |
| --- | --- | --- | --- |
| Missense mutations | 270 | 714 | 223 |
| Frame shift deletions | 11 | 20 | 5 |
| Frame shift insertions | 9 | 7 | 3 |
| Nonsense mutations | 18 | 54 | 16 |
